# Structure and Functions of Gesture Sequences in Wild Bonnet Macaques (*Macaca radiata*)

**DOI:** 10.1101/2024.03.18.585581

**Authors:** Shreejata Gupta, Anindya Sinha

## Abstract

Compositionality, a hallmark of human language, involves generating novel meaning by combining existing units. Nonhuman primates (mostly apes) are known to combine gestural units in non-random ways, but they do not make novel meaning with these combinations. What could, however, be the functional roles of these gesture sequences and whether they bear any significance to language evolution is still unclear. Moreover, studies on gesture-sequences in non-ape primate species is almost non-existent. Here, we investigated for the first time, the structure and functions of gesture sequences in the naturally occurring communication of wild bonnet macaques (*Macaca radiata)*, using analyses akin to ape gesture studies (Genty & Byrne, 2010; Hobaiter & Byrne, 2011). Bonnet macaque gesture sequences exhibit non-random combinations of gestures and non-gesture units – certain gestures are significantly more common in sequences, they associate preferentially with specific other components and certain components are more likely to appear either at the beginning or at the end of a sequence. Interplay of these sequences form distinct gestural clusters, corresponding to affiliative/play and agonistic contexts. Although, the overall functions of bonnet macaque gesture sequences remain obscure, as in apes, we found that gesture sequences were specifically used as a persistence strategy, after the initial single gestures have failed to initiate and sustain social interactions. We discuss our findings in the light of a possibility that primate gesture sequences, coordinating the flow of social interactions, may be evolutionary precursors to pragmatic gestures in human language.

## 1. Introduction

Compositionality is a hallmark of human language. We effortlessly recombine already meaningful words, gestures and facial expressions to compose novel meanings in the same or different contexts (Boer et al., 2012). Not only a feature of spoken language, compositionality is found across sign languages and other nonverbal communication in humans (e.g. facial expressions) (Cavicchio et al., 2018; Sandler & Lillo-Martin, 2006).

Evolutionary origins of this unique linguistic feature of compositionality remain elusive (Amici et al., 2022). Several studies have examined compositionality in primate gestures, suggested to lay the phylogenetic foundations of language (Corballis, 2002). Gesture sequences, composed of individually meaningful gestures in the species repertoire, have been long reported in primates, including in language-trained bonobos and chimpanzees (Greenfield and Savage-Rumbaugh, 1990, 1991; Brakke and Savage-Rumbaugh, 1995, 1996; Lyn et al., 2010), as well as in wild and zoo-living individuals (Plooij, 1978; Tomasello et al., 1994; Tomasello and Camaioni, 1997). These studies evoked ideas to test whether such gesture sequences in our closes phylogenetic relatives bear resemblances to syntactic constructions in human language (Tomasello et al., 1994).

Subsequent research challenged the notion of syntactic structure in ape gesture sequences. Captive chimpanzees frequently produce sequences of gestures, primarily consisting of repetitive tactile gestures, likely aimed at enhancing recipients’ responsiveness (Liebal et al., 2004). Similarly, Sumatran orangutans typically produce gesture sequences characterized by repeated gestures, with observations suggesting that these sequences may be largely driven by emotional arousal, irrespective of recipient response (Tempelmann & Liebal, 2012). Although it has been established that the structuring of these sequences in chimpanzees (Hobaiter & Byrne 2011) and in gorillas (Genty & Byrne, 2010) are not a random assembly of signal units, a clear idea about their functions in primate social interactions is lacking. While gorillas did not appear to use sequences to alter the meaning of communication, chimpanzees exhibited a gradual decrease in the frequency of gesture sequences with age, suggesting a developmental process akin to “repertoire-tuning” (Hobaiter and Byrne, 2011).

From a comparative perspective, studies on sequential use of gestures and their functions in non-ape species is rare. Aychet et al, (2021) conducted a detailed analyses of multimodal signal combinations (body, facial and vocal signals) in captive red-capped mangabeys, yet the authors, remain agnostic about the functions of these combinations. In this paper, we advance the state of this knowledge by providing evidence of gesture sequences in naturally occurring communication of a non-ape species (bonnet macaques, *Macaca radiata)*, and investigate the nature (structure and function) of these compositions. Bonnet macaques, an Old World Monkey species from southern India, possess a rich gestural repertoire (Gupta & Sinha 2019, 2016); they produce these gestures intentionally and flexibly, even as sequences with vocalisations (Deshpande et al., 2018).

In this paper, we analyse the gesture sequences in bonnet macaques, using Markov-transition analyses, previously used for ape-gesture sequences (Genty & Byrne, 2010; Hobaiter & Byrne, 2011), we determined the structure and function of these macaque gesture sequences, and how to compare to those found in ape gestural communication.

## 2. Methods

### 2.1. Study area

We conducted this study from February 2013 to July 2014 in the Bandipur National Park (11.66 °N, 76.63 °E) in the southern state of Karnataka, India. The most common primate species in this area is the bonnet macaque, the study species, described below. The troops that we studied belong to a free-ranging population, though leading a somewhat provisioned life along a highway that runs through the Park.

### 2.2. Study species

The bonnet macaque is a cercopithecine primate, endemic to peninsular India and extensively distributed across a wide range of habitats, possibly due to its exceptional ecological flexibility and behavioural lability, as has been documented in earlier studies (Sinha, 2001; Sinha et al., 2005; Ram et al., 2003). An elaborate behavioural repertoire and complex social interactions also characterize the species (Sinha, 2001). The gestural repertoire of this species comprises of 32 unique gestures (Gupta & Sinha., 2019).

For this study, SG identified four study troops in and around the Bandipur National Park, three of which had a species-typical multimale-multifemale social organization (Sinha, 2001), while one of them was a unimale-multifemale troop, an unusual, recently characterized form of social organization shown typically by this particular population (Dutta-Roy and Sinha, 2001; Sinha, 2001; Sinha et al., 2005, 2003). The study troops comprised a total of 29 adult females (>4 years of age, mean ± SEM = 7.25 ± 0.48), 23 adult males (>4 years of age, mean ± SEM = 5.75 ± 1.89), 31 juveniles (2–4 years of age, mean ± SEM = 7.75 ± 2.05) and 26 infants (0–2 years of age, mean ± SEM = 6.5 ± 1.04). Infants are entirely dependent on their mothers, especially during foraging and travelling. From a year after birth until their testicles have descended (for males) or they start cycling (for females) is considered the juvenile stage, after which individuals are considered adults.

### 2.3. Data coding

SG conducted focal animal sampling (Altmann, 1974) to manually documented gestures in the study population following the published repertoire for this species (Gupta & Sinha 2019). SG also collected adlib video recordings at the initial stage of the study used to assess inter-observer reliability by a second researcher, who was not a part of this particular study, but has been exposed to macaque and ape behaviour studies as part of her own research. The randomly chosen videos covered 81% of the reported gestures and the Cohen’s Kappa value for all the gestures ranged from 0.80 to 1.00 (except the gesture Touch, for which the reliability score was 0.56).

Following the definitions used by Genty et al, (2010) and Hobaiter & Byrne (2011), we defined a gesture sequence as a series of gestural acts displayed one after the other within an interval of <1 second. Moreover, we documented several sequences where gestures were associated with other non-gesture actions and body postures. We also consider these compositions (gesture-other compositions) here and analyse them separately. Data was pooled across all individuals in the four study troops for the present analysis.

### 2.4. Data analyses

The sequence lengths and their frequency of use in various contexts were compared with those of single gestures, adopted from Liebal et al, (2004) and Genty and Byrne, (2010). We also examined the probability of certain gestures being used more often singly or as components of sequences.

In order to understand the structure of the sequences and determine the probability of transition from one gesture / non-gesture to the others, we applied Markov-transition analyses, described in Genty & Byrne (2010). A matrix of pairwise transitions was formed with the corresponding frequencies of transition from one gesture / non-gesture to the others (Fabricius & Jansson 1963) and the transition which occurred at least four times during the entire study was considered for further analysis (see Genty & Byrne 2010). The probabilities of any two gesture / non-gesture composition were then tested for their significance of association using the binomial test in the software R (version 4.3.1, 2023-06-16). We also investigated whether there were certain gestures that initiated or ended a sequence at significantly higher probabilities than did others.

To determine the functions of gesture sequences, we carried out the following analyses (see Genty & Byrne 2010):

1. a comparison between the relative success (measured as elicitation of response in receivers) of repeated gestures in a sequence and that of this gesture used singly;
2. a comparison of the relative success of gesture / non-gesture sequences than that of the combined effects of each component performed alone;
3. preference of signallers to use gesture sequences in case of persistence in communication, in the initial absence of a response, over using the components alone.

In these three sets of analyses, we evaluated the success of the test gestures or sequences by measuring the proportion of times appropriate responses were elicited in relation to those in which they were not (‘no-response situations’). Our analysis of the second point listed above, involved only the sets of two-component-long sequences. We first separately calculated the probability of elicitation of no-responses for each of the two components of these sequences, [for example, p(A) and p(B) of two particular components A and B respectively] and then computed the expected combined probability of no-response when both were displayed independently by multiplying their individual probabilities [p(A) × p(B)]. This value was then compared with the observed probability of no-response when these components were performed in a sequence [p(A-B)]. We hypothesise that if p(A) × p(B) is greater than p(A-B) the sequence, as a whole, is more effective in evoking a response than when the two components are exhibited in isolation (see Genty and Byrne 2012).

## 3. Results

We found two kinds of compositions possible for gesture sequences in bonnet macaques— gesture-gesture compositions and gesture-other (with the non-gestures) compositions; both of these associations were analysed independently. All the 32 gestures in the bonnet macaque repertoire occurred in these observed sequences (Appendix 1, Table A1), while there were 16 non-gestures—Bared-Teeth Displaying, Branch-Shaking, Copulatory-Bobbing Vertically, Fear-Grimacing, Gazing, Ground-Slapping, Inspecting by Smelling, Inspecting by Tasting Oestrous Material, Inspecting by Touching, Inspecting Visually, Leaping Away, Licking, Mounting, Mounting with Lip-Smacking, Sniffing and Touching Nipples—that featured in the gesture-other compositions (Appendix 1, Table A2). A total of 3283 gesture tokens were analysed, comprising of 72.13% single gestures (2368 tokens) and 27.87% gesture sequences (915 tokens).

### 3.1. Sequence length and context of use

The frequencies of occurrence of all the observed sequences were compared with that of single gestures or non-gestures. Single gestures were displayed more frequently by the study individuals than were the sequences (**Table 1**). Of the 915 observed sequences, 812 were gesture-gesture compositions, while 64 2-length, 23 3-length and 16 4-length sequences were gesture-other compositions. The comparison of the contexts of use revealed significantly more use of single gestures in the contexts of affiliation and play (Affiliation: G-test of independence, df = 1, G = 372.73, *p* < 0.001, n = 1446, Play: df = 1, G = 75.45, *p* <0.001, n = 403; **Figure 1**). There were no such differences in the agonistic contexts.

**Table 1:** Number of gesture sequences of various lengths displayed by bonnet macaques in the four study troops.

| Number of components | Total |
| --- | --- |
| 2 | 722 |
| 3 | 130 |
| 4 | 46 |
| 5 | 11 |
| >5 | 6 |
| **Total number of sequences** | **915** |

**Figure 1:**
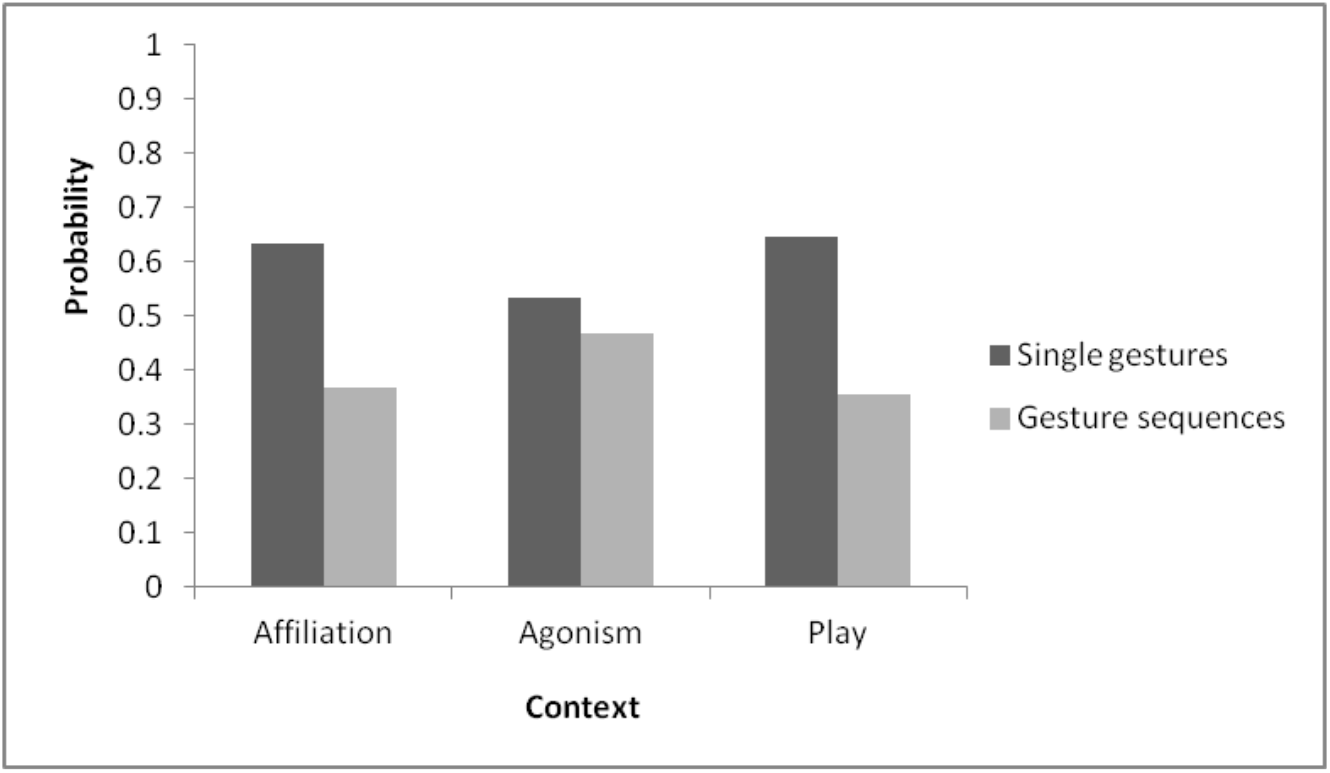
Probability of gestures and gesture sequences in different contexts, in bonnet macaque communication

We evaluated whether all the 32 gestures and the 16 non-gestures were comparably used by the study macaques as independent units or as components of sequences (**Figure 2**). Bared-Teeth Displaying and Copulatory-Bobbing Vertically were never used singly and hence, removed from this analysis. The results revealed 12 gestures and 3 non-gestures to be significantly preferentially used in sequences while 13 gestures and one non-gesture were more likely to be used as independent units, in this population (**Table 2**).

**Table 2:**
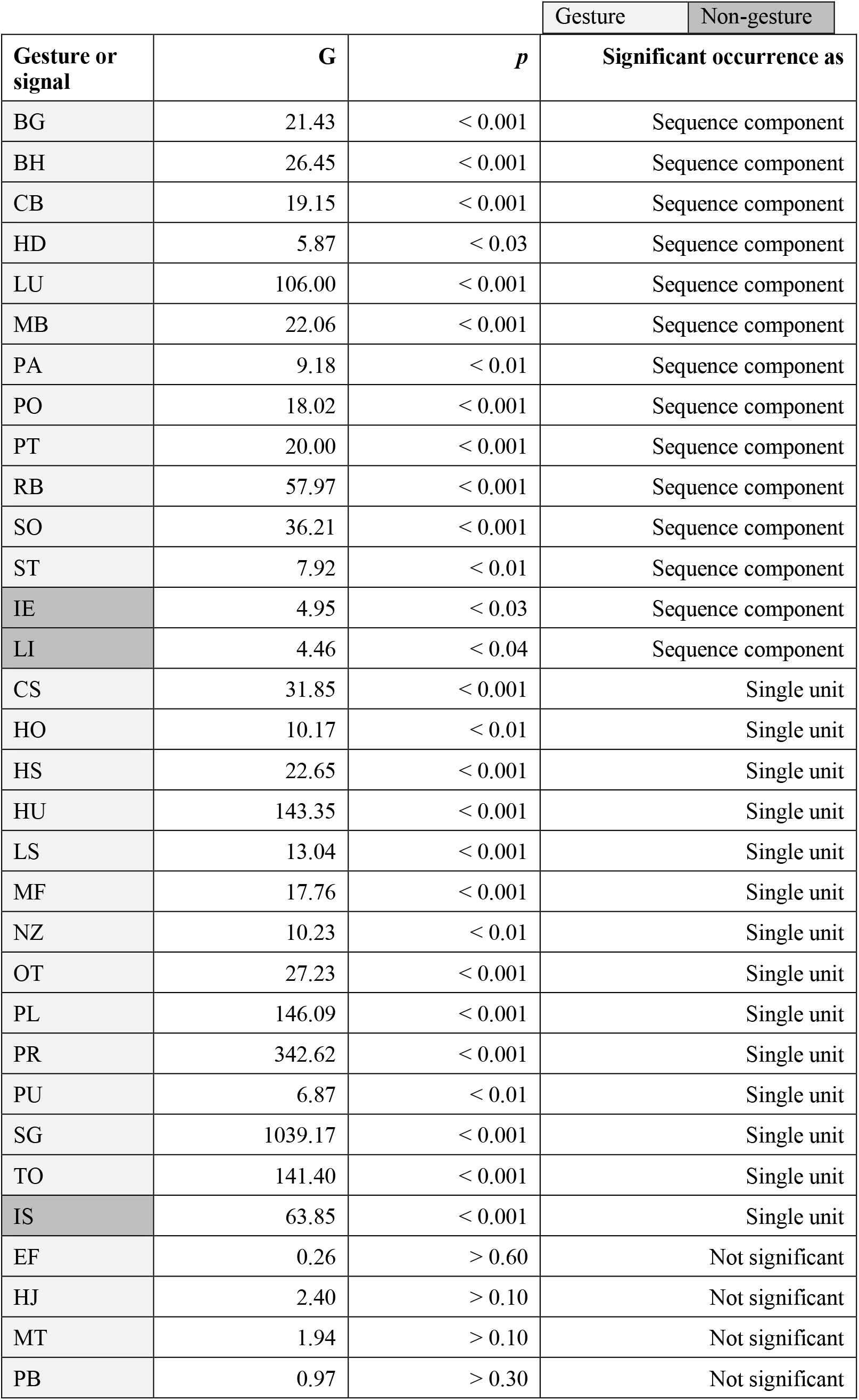
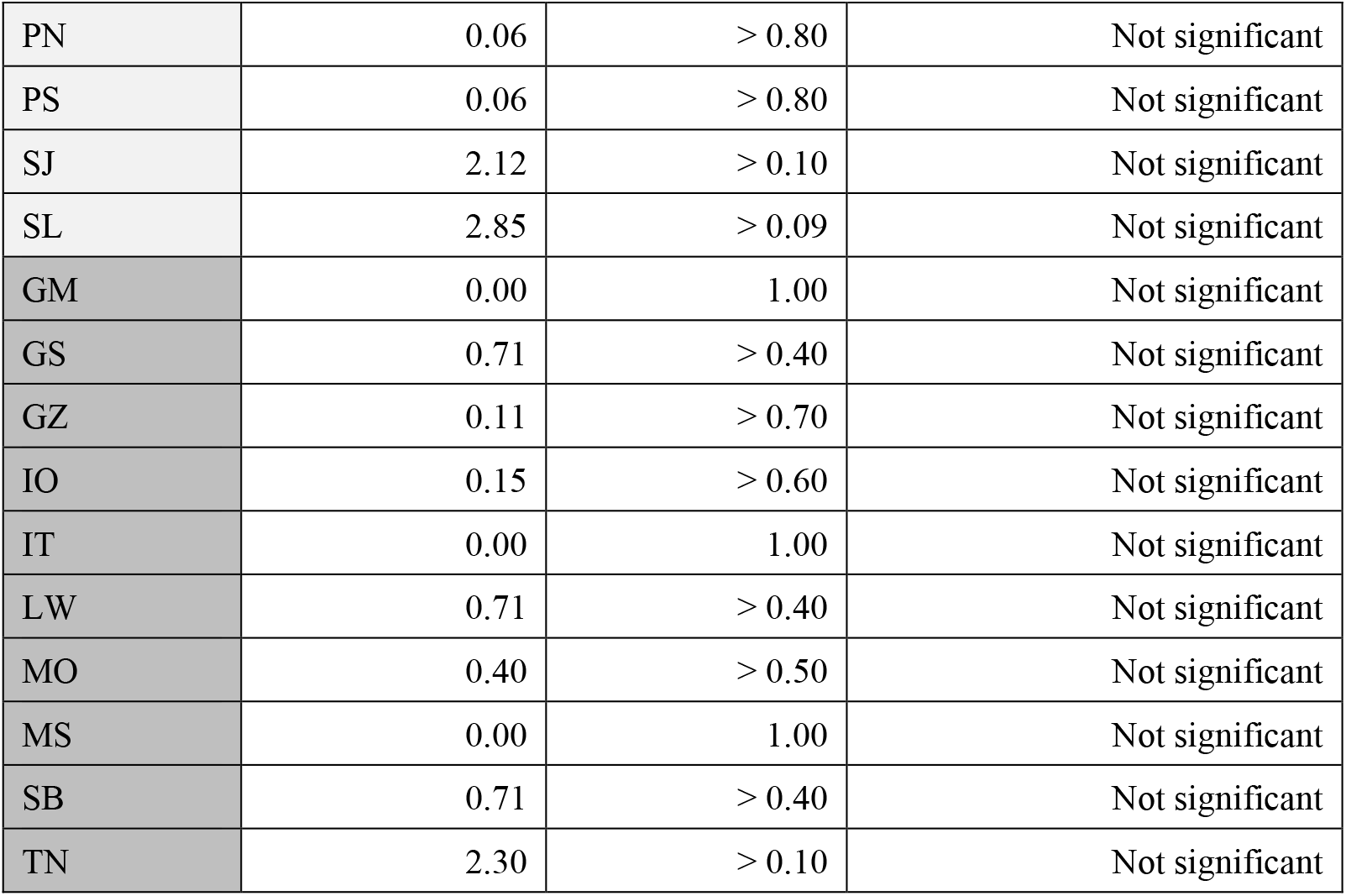
Gestures and non-gestures that were significantly used either as single unit or in a sequence by bonnet macaques. Proportions were compared using the G-test of independence.

**Figure 2:**
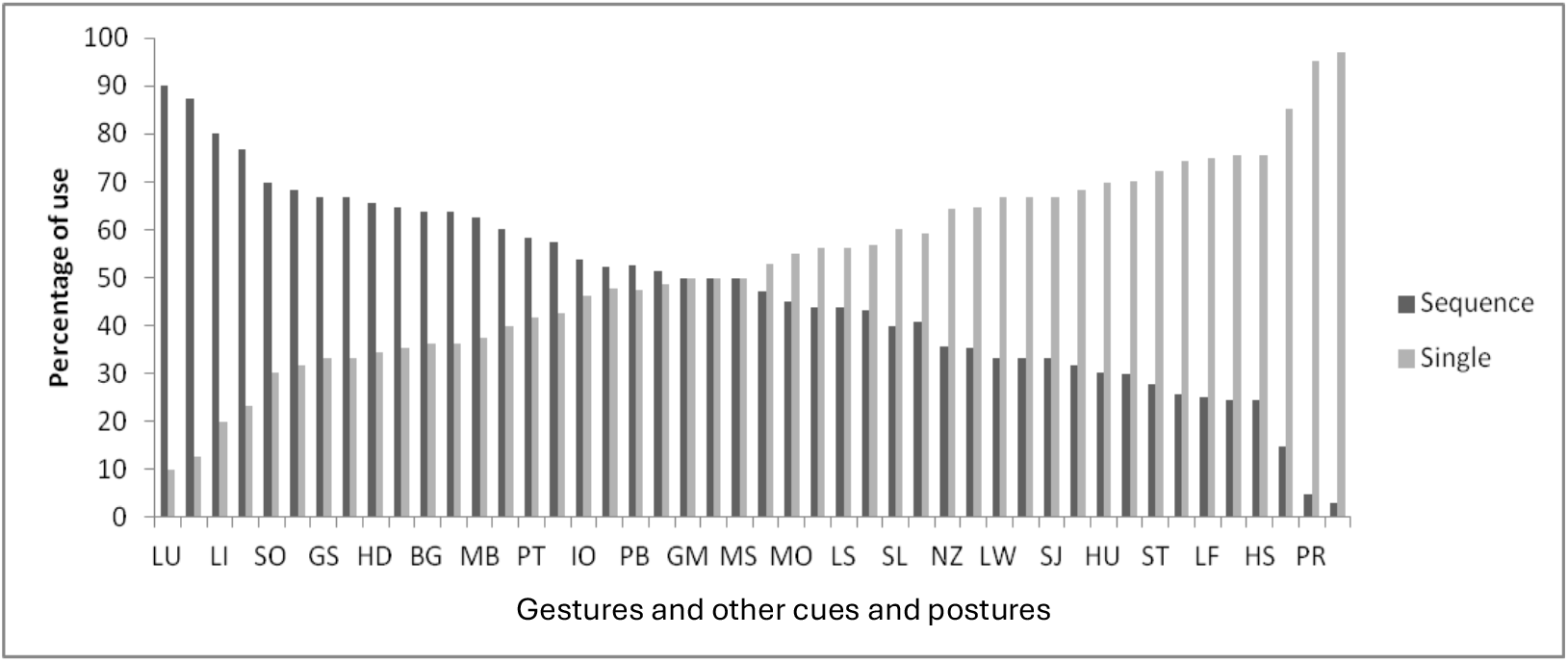
Proportional use of gestures and non-gestures as single units or in sequences, by bonnet macaques

### 3.2. Structure of gesture sequences

The structure of the gesture sequences was analysed by Markov-transition method, in order to test significant associations between any two gestures or non-gestures. There were 372 gesture-gesture and gesture-other dyads, of which only 99 of them occurred at least four times during the observation. Of these 99 dyads, 33 gesture-gesture compositions occurred significantly more than others and four gesture-other compositions were significantly paired. Among the 33 gesture dyads, seven were repetitions of the same gestures, composed of Biting Gently, Head-Jerking, Mouth-to-Mouth Sniffing, Patting, Pulling, Raising Eyebrows and Touching. The rest 26 gesture-gesture dyads formed two main clusters, one with affiliative and play gestures (**Figure 3a**), the other with agonistic gestures (**Figure 3b**). The four gesture-other compositions have been presented separately (**Figure 3c**).

**Figure 3:**
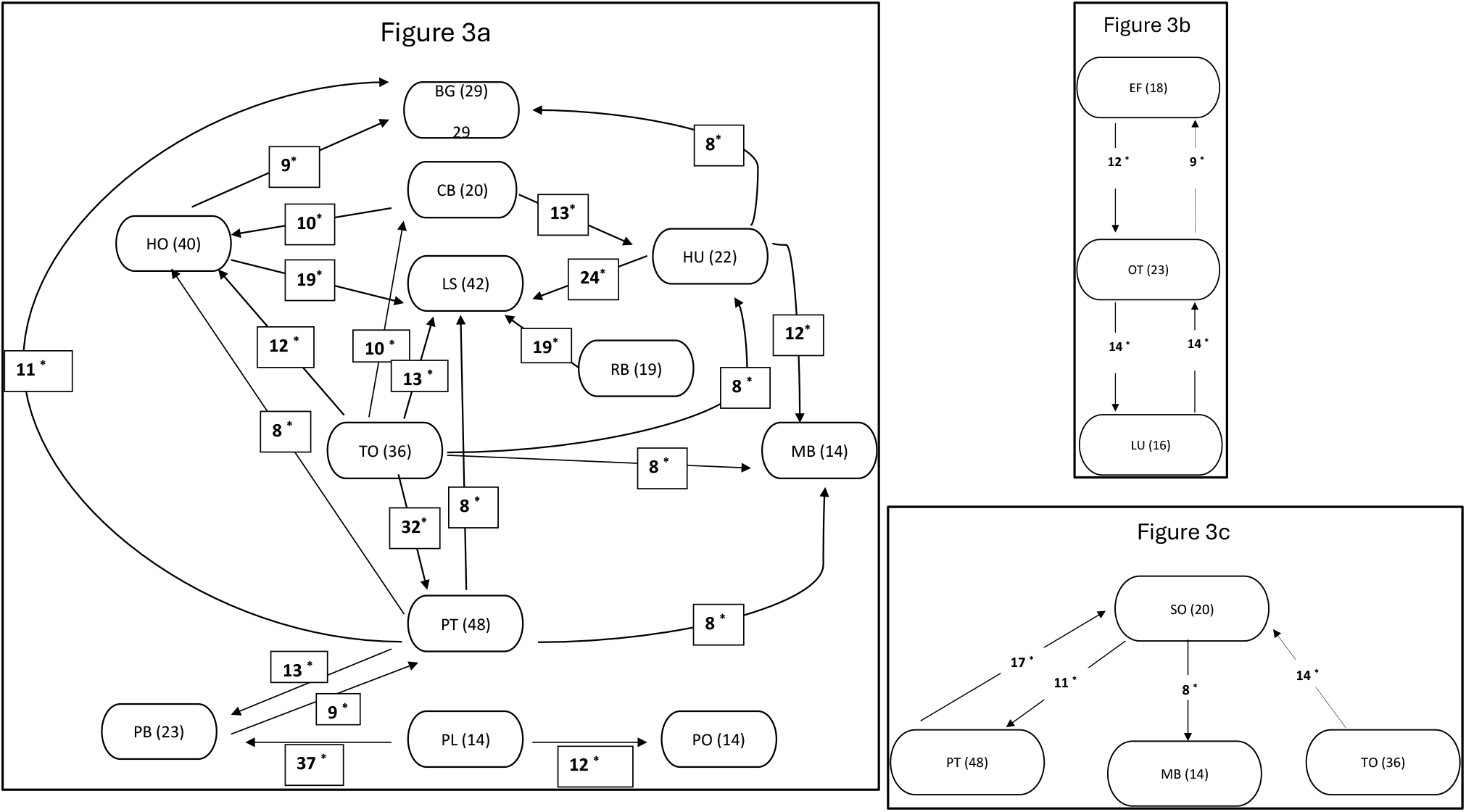
Network of gesture-gesture compositions in (a) affiliative and play and (b) agonistic contexts; (c) gesture-other compositions, in bonnet macaques. Each box represents the gestures (Table 2) along with the number of sequences they occurred in. The arrows indicate the gesture significantly associated with the preceding one. The numbers next to the arrows signify the occurrence frequency of each gesture-gesture pair. The asterisks depict significant associations of the components, as evaluated by the binomial test

There were nine particular gestures that were used at the beginning of a sequence and ten at the end of a sequence with significantly higher probabilities than were the other components (**Table 3, 4**).

**Table 3:**
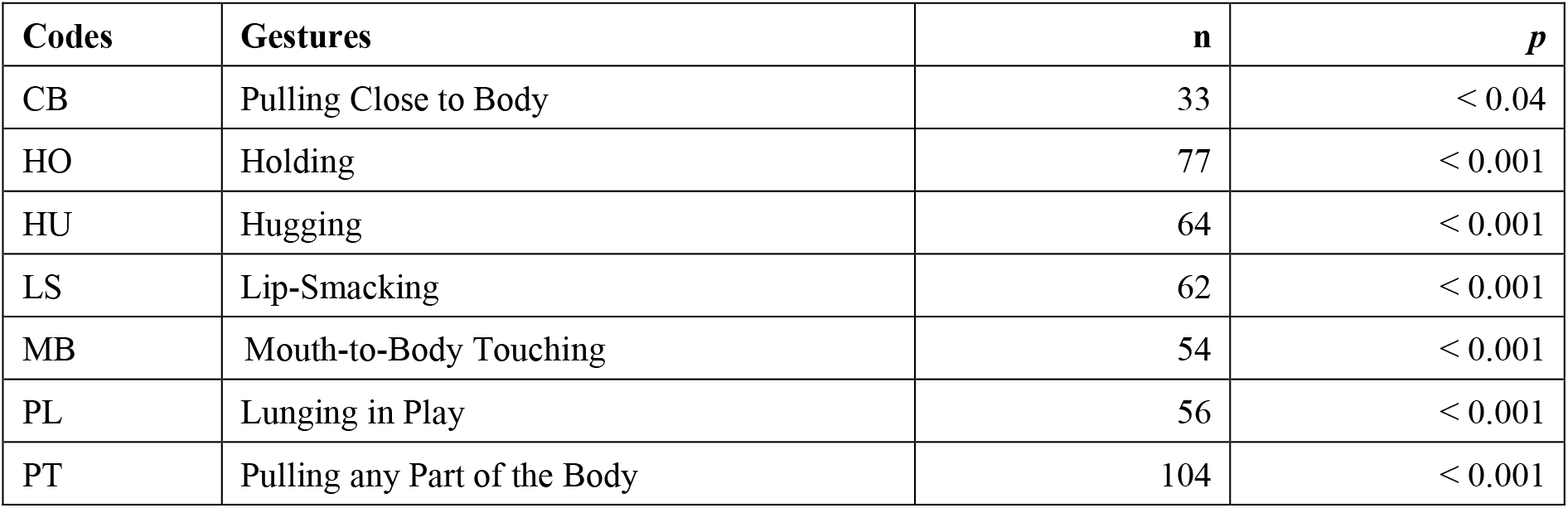
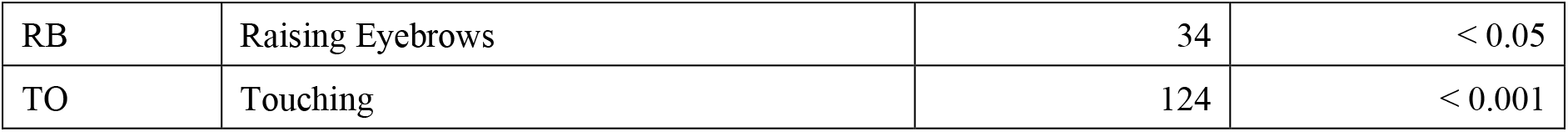
Gestures used significantly more at the beginning of a sequence, in bonnet macaques( evaluated by the binomial test)

**Table 4:** Gestures or non-gestures used significantly more at the end of a sequence, by bonnet macaques (evaluated by the binomial test)

| Codes | Gestures or Signals | n | p |
| --- | --- | --- | --- |
| BG | Biting Gently | 64 | < 0.001 |
| HO | Holding | 60 | < 0.001 |
| HU | Hugging | 58 | < 0.001 |
| LS | Lip-Smacking | 106 | < 0.001 |
| MB | Mouth-to-Body Touching | 62 | < 0.001 |
| OT | Open-Mouth Threatening | 41 | < 0.001 |
| EB | Biting in Play | 69 | < 0.001 |
| PT | Pulling any Part of the Body | 95 | < 0.001 |
| TO | Touching | 37 | < 0.001 |
| SO | Sniffing * | 33 | < 0.01 |

\* Sniffing is a non-gesture signal

### 3.3. Gesture repetitions, sequences and their functions

First, we compared the relative success of gesture repetitions with that of the component gesture used singly. Success was measured in terms of the proportion of times they elicited an appropriate response in the receiver, as compared to those that did not (‘no-response situations’). The proportions of appropriate responses elicited by the gesture repetitions (23 of 48 events) and their component gestures used singly (345 of 659 events) were not significantly different (G = 0.63, df = 1, *p* > 0.40) (**Figure 4**). We also did not find any significant relation between the length of these repeat-sequences to the probability of an elicited responses in the target audience (Spearman’s rank correlation, rho = -0.26, n = 17, *p* > 0.20; **Figure 5**). This indicated that the sequential repetition of gestures did not function as an efficient strategy to elicit responses to continue social interactions from the target audience.

**Figure 4:**
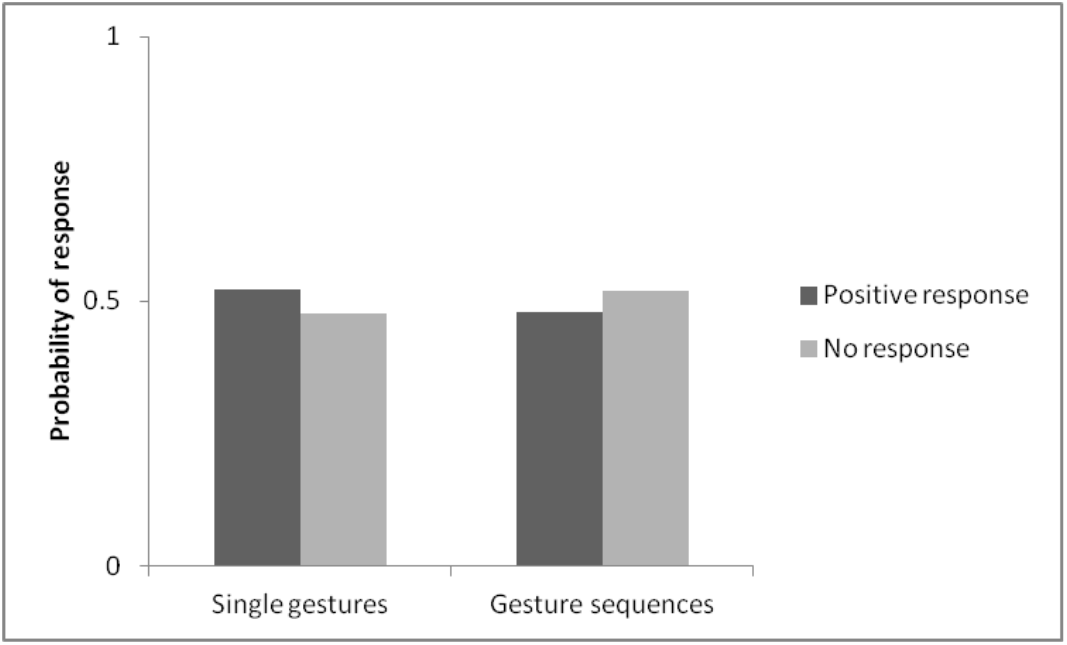
Probability of responses or no-responses elicited by single and repeated gestures in bonnet macaques

**Figure 5:**
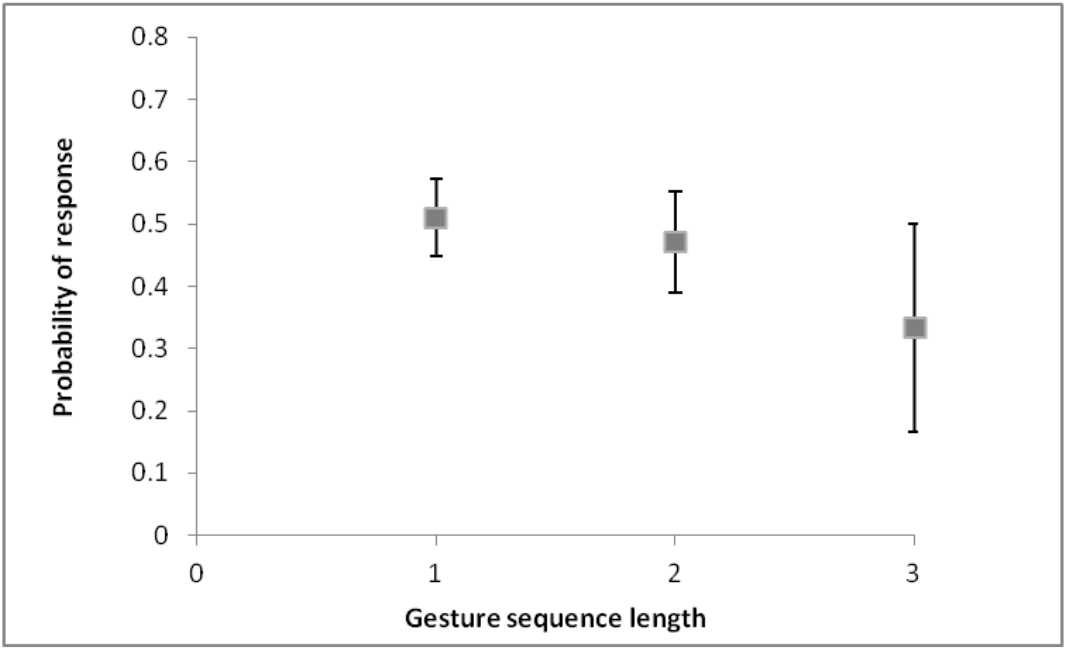
Probability of responses elicited by repeated gestures in a sequence of increasing lengths, in bonnet macaques

We also investigated whether gesture sequences, consisting of two distinct components, were relatively more effective in eliciting responses than when each of these components were performed singly. The 33 gesture-gesture dyads and the four gesture-other dyads that had significantly greater probabilities of association were considered in this analysis. The observed probabilities of no-response were found to be higher for 26 of the 29 two-component gesture sequences, indicating that these sequences were not more effective in eliciting a response than were their components performed independently. Detail list of the probabilities of each of these pairs are provided in Appendix 1, Table A3.

### 3.4. Strategies of persistent gesturing

We evaluated whether signallers prefer to use gesture sequences to display persistence, in the initial absence of a response, instead of persisting with single gestures.

Single gestures were displayed 2368 times during the observation period, of which 899 instances (37.96%) constituted no-response situations. In these situations, the signalling individual terminated the communication process in 807 instances (89.77%) while in the remaining 92 instances (10.23%), the signaller persisted until the intended goal was achieved. For persistent gesturing, bonnet macaques used gestures sequences in 26.44% (23 out of 92) of these cases, the repeated the original gesture singly in 48.28% (42 out of 92) and alternative, but functionally similar, gestures used singly on in 25.29% (27 out of 92) occasions. Assuming that the study individuals were equally likely to use any of these strategies to persistently communicate with the recipient, they prefer to repeat the initial gesture singly, following the failure of the recipient to respond in the first instance (G-test of independence, G = 18.50, df = 2, *p* < 0.001).

But were the persistent displays of the initially used single gesture more successful in eliciting responses in the targetted recipients? **Table 5** depicts the frequencies of appropriate and no-responses elicited in the receivers when the signallers used the three strategies—same gestures repeated singly, gestures in a sequence, and alternative, functionally similar, single gestures. A comparison of the effectiveness of these three strategies made it evident that the performance of both gesture sequences and alternative, functionally similar, single gestures were more successful than the same gestures repeated singly in evoking greater proportion of appropriate responses (Gesture sequences and same gestures: G = 11.84, df = 1, *p* < 0.001; Alternative gestures and same gestures: G = 5.65, df = 1, *p* < 0.02). Gesture sequences were, however, even more effective than the alternative single gestures in this regard (Gesture sequences and alternative gestures: G = 8.75, df = 1, *p* < 0.01).

**Table 5:** Frequencies of appropriate and no responses received in response to different strategies during occasions of persistent gesturing by the study bonnet macaques.

| Strategy | Appropriate response | No response |
| --- | --- | --- |
| Gesture sequence | 17 | 6 |
| Same gesture, repeated singly | 22 | 20 |
| Alternative, functionally similar, single gesture | 17 | 10 |

## 4. Discussion

Bonnet macaques use two kinds of gesture sequences – gesture-gesture combinations and gesture-other (other non-gesture actions and body postures) combinations of varying lengths. The non-gesture components were not considered as gestures as, there were insufficient number of occurrences of some these units (Bared-Teeth Displaying, Copulatory-Bobbing Vertically and Gazing) in the population to test their the criteria of qualifying as intentional gestures. Moreover, for the 13 other non-gestures— Branch-Shaking, Fear-Grimacing, Ground-Slapping, Inspecting by Smelling, Inspecting by Tasting Oestrous Material, Inspecting by Touching, Inspecting Visually, Leaping Away, Licking, Mounting, Mounting with Lip-Smacking, Sniffing and Touching Nipples—the joint attention state between the signaller and recipient did not prevail in most of the events when the signaller displayed these particular signals and they could not, therefore, be defined as true gestures. It is, however, noteworthy that these signals were used in conjunction with other gestures, incorporated into gestural sequences and, thus, formed an integrative communication system with gestures. It is possible that these actions and postures could ultimately qualify as gestures with additional observations and recorded instances.

### 4.1. Length of sequences and their contexts of use

The lengths of the gesture sequences produced by bonnet macaques varied from two to five components and were also less frequently produced than were single gestures, similar to reports from gorillas and chimpanzees (Genty and Byrne 2010; Hobaiter and Byrne 2011). Affiliative and play gestures were used significantly more as single units than in sequences by the macaques, contrasting to what has been observed in chimpanzees and gorillas, wherein the most frequent sequences consisted of play gestures (Liebal et al. 2004; Genty and Byrne 2010; Hobaiter and Byrne 2011). It, thus, appears that bonnet macaques prefer to communicate through single gestures although a certain proportion of their communication is indeed represented by sequential combinations. Certain gestures in the bonnet macaque repertoire were significantly more used in sequences than as single units and vice versa. This could be an indication that certain gestures are sufficient by themselves in information transfer, while others are more effective in combination with other components.

### 4.2. Structure of the gesture sequences

The Markov-transition analysis revealed that the gesture sequences in bonnet macaques did indeed have above chance preferential associations between gestures and non-gestures, while others had high probabilities of being repeated sequentially. More remarkably, certain components of these sequences were more likely to be incorporated at the beginning or at the end of a sequence than were others. Similar observations have been made earlier for gorillas, wherein certain gestures formed a network with other gestures, at probabilities higher than would be expected by chance (Genty and Byrne 2010).

In our study, we found two distinct structural clusters made of significantly associated gestures in each of them. It is illuminating that one of these networks consisted of affiliative and play gestures while agonistic gestures alone constituted a separate cluster. What must be noted here is that these two structural clusters of closely associated gestures appeared to be functionally distinctive and, therefore, possibly served very different roles bonnet macaque communication. Could these contextually related gesture clusters be compared to semantic-maps of words? Future analyses in these lines could be interesting in relation to language evolution. We also detected a third network consisting of a few gestures and non-gestures, including Pulling any Part of the Body (PT), Sniffing (SO), Touching (TO) and Mouth-to-Body Touching (MB), the functional role of which was not very evident but which could serve either as an affiliative or a curiosity-driven sequence of behavioural interactions, occasionally leading to sexual inspection.

It may be relevant to mention here that the gestures—Pulling any Part of the Body (PT) and Touching (TO)—were preferentially associated with several other gestures and non-gestures with very high probabilities of occurrence. These two gestures were also often found at the beginning of the sequences. Interestingly, we earlier reported that it was hard or impossible to determine the function of these two particular gestures (Gupta & Sinha 2019) and we hypothesised that perhaps these gestures were more often used in association with other gestures, for which we find evidence in our present study. Tactile gestures like PT and TO could be acting as “attention-getters” (Tomasello et al. 1994) and could invariably be performed before other gestures, which actually conveyed the true intention of the communication event once the attention of the recipients were secured.

We also speculate that another possible function of these two observed gestures could be to leave open-ended opportunities to constantly manipulate the communication sequence to their advantage, either navigating it towards affiliative behaviours, or towards play (Figure 3a), similar in principle to what has also been suggested by Genty and Byrne (2010). Future studies on gesture sequence analysis, incorporating signals from other modalities should be able to shed light on these speculated functions.

### 4.3. Gesture repetitions and sequences: Advantages over single gestures?

The length of the repeated-gesture sequences did not appear to significantly increase communication success, in terms of eliciting response in the target audience. This is contrary to what is reported for repeat-sequences of gestures in captive chimpanzees (Liebal et al., 2004) and Sumatran orangutans (Tempelmann & Liebal, 2012). Moreover, for the overwhelming majority of two-component gesture sequences, the sequences themselves did not seem to elicit significantly more responses from the audience, as compared to the combined effect of the same two gestures performed singly. Thus, the overall function of gesture sequences in bonnet macaques remain obscure, similar to what has been concluded earlier for the gesture sequences produced by wild gorilla populations (Genty and Byrne 2010).

### 4.4. Strategies of persistent gesturing

When an initial single gesture failed to evoke any response from the targeted receiver, bonnet macaques preferred to repeat the initial, single gesture rather than taking refuge into gesture sequences. In these special situations of persistence, however, the gesture sequences were functionally most effective in evoking a response and continue the social interaction, than single gesture units. Do gesture sequences in bonnet macaques have pragmatic functions acting as interactional gestures? Are primate gesture sequences, in general, the pragmatic drivers of interaction, like some human co-speech gestures (Bavelas et al., 1992), rather than generating novel semantic functions as a result of compositionality? Taken together with the roles of certain tactile gestures in bonnet macaques offering fluidity to sequences, allowing flexibility in the flow of interactions at will (as discussed above), the pragmatic roles of gesture sequences in primates presents a promising scope of future research.

### 4.5. Limitations

In this study, we did not consider multimodal signals and their interactions with gestures in these sequences. Our focus was limited only to visual and tactile bodily signals, ignoring facial expressions and vocalisations. Similar holistic analyses as performed by Aychet et al, (2021), with more focus on functional roles of such multimodal, multicomponent sequences, should pave the future way in the domain of compositionality research in primate communication. In this context, one should also investigate the temporal alignment of different components in sequences and apply conversational analysis methods, as done in human language, to truly unravel the evolutionary origins of compositionality in language. We also could not integrate a developmental perspective in this present study due to insufficient documentation of sequences across age. Future studies tracing the ontogeny of these sequence use in bonnet macaques, and other primate species, will provide a better understanding of their functions, as pointed out by Hobaiter and Byrne (2011).

## Acknowledgements

This study, carried out within the National Institute of Advanced Studies, Bangalore, India, has benefitted from the support of the research grant under the Cognitive Science Research Initiative (Grant Number SR/CSI/44/2008) awarded to AS and a doctoral fellowship from the National Institute of Advanced Studies, awarded to SG. Both authors are grateful to Kantharaju H C of the Karnataka Forest Department for granting permission for this work, Jagadish M and Sharmi Sen for help during data collection, Kate Grounds for coding video for inter-observer reliability. Karthik Davey and Sukanta Das had been instrumental for this study, providing logistical support during fieldwork.

## Appendix 1

**Table 1:** Component gestures displayed in sequences by bonnet macaques.

| Code | Gesture | Context |
| --- | --- | --- |
| LS | Lip-Smacking | Affiliation |
| SG | Soliciting Allogrooming | Affiliation |
| BG | Biting Gently | Affiliation |
| CB | Pulling Close to Body | Affiliation |
| HO | Holding Gently | Affiliation |
| HS | Hugging with Lip-Smacking | Affiliation |
| HU | Hugging | Affiliation |
| MB | Mouth-to-Body Touching | Affiliation |
| MT | Mouth-to-Mouth Touching | Affiliation |
| NZ | Nuzzling | Affiliation |
| PA | Patting | Affiliation |
| TO | Touching | Affiliation |
| HJ | Head-Jerking | Agonism |
| LU | Lunging | Agonism |
| OT | Open-Mouth Threatening | Agonism |
| BH | Biting Hard | Agonism |
| EF | Eye Flashing | Agonism |
| HD | Holding Down Roughly | Agonism |
| SL | Slapping | Agonism |
| PL | Lunging in Play | Play |
| PO | Open-Mouth Threatening in Play | Play |
| SJ | Spot-Jumping | Play |
| PB | Biting in Play | Play |
| PS | Slapping in Play | Play |
| MO | Mounting | Multifunctional |
| PR | Presenting | Multifunctional |
| PT | Pulling any Part of the Body | Multifunctional |
| PU | Pushing | Multifunctional |
| PN | Pinching | Multifunctional |
| ST | Staring | Multifunctional |
| RB | Raising Eyebrows | Multifunctional |
| CS | Copulatory Lip-Smacking | Multifunctional |
| **Total number of gestures = 32** |  |  |

**Table 2:** Component signals displayed in sequences by bonnet macaques.

| **Code** | **Gesture** | **Context** |
| --- | --- | --- |
| BT | Bared-Teeth Displaying | Agonism |
| GM | Fear-Grimacing | Agonism |
| GS | Ground-Slapping | Agonism |
| GZ | Gazing | Sexual |
| IS | Inspecting by Smelling | Sexual |
| IE | Inspecting by Tasting Oestrous Material | Sexual |
| IT | Inspecting by Touching | Sexual |
| IO | Inspecting Visually | Sexual |
| LW | Leaping Away | Affiliation |
| LI | Licking | Affiliation |
| MO | Mounting | Dominance-subordination |
| MS | Mounting with Lip-Smacking | Dominance-subordination |
| SB | Branch-Shaking | Affiliation |
| SO | Sniffing | Agonistic |
| TN | Touching Nipples | Neutral |
| UB | Copulatory-Bobbing Vertically | Sexual |
| **Total number of signals = 16** |  |  |

**Table 3:** Comparison of the calculated expected and observed probabilities of no-responses elicited by two gestures or signals, displayed independently and in two-component gesture sequences respectively.

| Sequence | n | Expected probability | Observed probability |
| --- | --- | --- | --- |
| CB-HO | 10 | 0.17 | 0.43 |
| CB-HU | 13 | 0.09 | 0.22 |
| EF-OT | 12 | 0.14 | 0.44 |
| HO-BG | 9 | 0.25 | 0.83 |
| HO-LS | 19 | 0.21 | 0.4 |
| **HU-LS** | **24** | **0.11** | **0.10** |
| HU-BG | 8 | 0.13 | 0.43 |
| HU-MB | 12 | 0.19 | 0.67 |
| **LU-OT** | **14** | **0.05** | **0.00** |
| OT-EF | 9 | 0.14 | 0.5 |
| **OT-LU** | **14** | **0.05** | **0.00** |
| PB-PT | 9 | 0.09 | 0.33 |
| PL-PB | 37 | 0.04 | 0.29 |
| PL-PO | 12 | 0.03 | 0.57 |
| PT-HO | 8 | 0.25 | 0.50 |
| PT-LS | 8 | 0.18 | 1.00 |
| PT-BG | 11 | 0.21 | 0.60 |
| PT-PB | 13 | 0.09 | 0.50 |
| PT-MB | 8 | 0.31 | 1.00 |
| PT-SO | 17 | 0.31 | 0.67 |
| RB-LS | 19 | 0.34 | 0.75 |
| SO-PT | 11 | 0.31 | 0.80 |
| TO-HO | 12 | 0.25 | 0.78 |
| TO-LS | 13 | 0.18 | 0.50 |
| TO-PT | 32 | 0.21 | 0.76 |
| TO-HU | 8 | 0.13 | 0.50 |
| TO-CB | 10 | 0.15 | 0.33 |
| TO-MB | 8 | 0.31 | 0.50 |
| TO-SO | 14 | 0.31 | 0.50 |

Cases where the observed probability was less than that of the expected probability have been highlighted in bold

## References

1. Altmann, J. (1974). Observational study of behavior: sampling methods. Behaviour, 49(3-4), 227–266.

2. Amici, F., Oña, L., & Liebal, K. (2022). Compositionality in primate gestural communication and multicomponent signal displays. International Journal of Primatology, 1–19.

3. Aychet, J., Blois-Heulin, C., & Lemasson, A. (2021). Sequential and network analyses to describe multiple signal use in captive mangabeys. Animal Behaviour, 182, 203–226.

4. Brakke K E and Savage-Rumbaugh E S. 1995. The development of language skills in bonobos and chimpanzee – 1. Comprehension. Language and Communication 13: 315–338.

5. Brakke K E and Savage-Rumbaugh E S. 1996. The development of language skills in Pan – II. Production. Language and Communication 16: 361–380.

6. Cavicchio, F., Dachkovsky, S., Leemor, L., Shamay-Tsoory, S., & Sandler, W. (2018). Compositionality in the language of emotion. PloS one, 13(8), e0201970.

7. Corballis, M. C. (2002). From hand to mouth: The origins of language. Princeton University Press.

8. De Boer, B., & Verhoef, T. (2012). Language dynamics in structured form and meaning spaces. Advances in Complex Systems, 15(03n04), 1150021.

9. Deshpande, A., Gupta, S., & Sinha, A. (2018). Intentional communication between wild bonnet macaques and humans. Scientific reports, 8(1), 5147.

10. Fabricius, E., & Jansson, A. M. (1963). Laboratory observations on the reproductive behaviour of the pigeon (Columba livia) during the pre-incubation phase of the breeding cycle. Animal Behaviour, 11(4), 534–547.

11. Genty E and Byrne RW. 2010. Why do gorillas make sequences of gestures? Animal Cognition 13: 287–301.

12. Greenfield P M and Savage-Rumbaugh E S. 1990. Grammatical combination in Pan paniscus: Processes of learning and invention in the evolution and development of language. In Sue Taylor Parker K R G (Ed.), ‘Language’ and Intellignece in Monkeys and Apes: Comparative Developmental Perspectives, Cambridge, New York: Cambridge University Press, pp. 540–578.

13. Greenfield P M and Savage-Rumbaugh E S. 1991. Imitation, grammatical development and the invention of protogrammar by an ape. In fKrasengor N A, Rumbaugh D M, Schiefelbusch R L and Studdert-Kennedy M (Eds.), Biological and Behavioral Determinants of Language Development, Hillsdale, New Jersey, USA: Lawrence Erlbaum, pp. 235–258.

14. Gupta, S., & Sinha, A. (2016). Not here, there! Possible referential gesturing during allogrooming by wild bonnet macaques, Macaca radiata. Animal Cognition, 19, 1243–1248.

15. Gupta, S., & Sinha, A. (2019). Gestural communication of wild bonnet macaques in the Bandipur National Park, Southern India. Behavioural processes, 168, 103956.

16. Hobaiter C and Byrne R W. 2011. Serial gesturing by wild chimpanzees: Its nature and function for communication. Animal Cognition 14: 827–838.

17. Liebal K, Call J and Tomasello M. 2004. Use of gesture sequences in chimpanzees. American Journal of Primatology 64: 377–396.

18. Lyn H, Greenfield P M and Savage-Rumbaugh E S. 2010. Semiotic combinations in Pan: A comparison of communication in a chimpanzee and two bonobos. First Language 31: 300–325.

19. Plooij F X. 1978. Some basic traits of language in wild chimpanzees? In Lock A (Ed.), Action, Gesture and Symbol: The emergence of Language, London UK: Academic Press, pp. 111–131.

20. R Core Team (2021). R: A language and environment for statistical computing. R Foundation for Statistical Computing, Vienna, Austria. Retrieved from https://www.R-project.org/

21. Sandler, W., & Lillo-Martin, D. C. (2006). Sign language and linguistic universals. Cambridge University Press.

22. Sinha, A. (2001). The bonnet macaque revisited: ecology, demography and behaviour. ENVIS Bulletin: Wildlife and Protected Areas, 1, 30–39.

23. Sinha, A. (2003). A beautiful mind: Attribution and intentionality in wild bonnet macaques. Current Science, 1021–1030.

24. Sinha, A. (2005). Not in their genes: Phenotypic flexibility, behavioural traditions and cultural evolution in wild bonnet macaques. Journal of biosciences, 30(1), 51–64.

25. Tempelmann, S., & Liebal, K. (2012). Spontaneous use of gesture sequences in orangutans. Developments in primate gesture research, 6, 7.

26. Tomasello, M., Call, J., Nagell, K., Olguin, R., & Carpenter, M. (1994). The learning and use of gestural signals by young chimpanzees: A trans-generational study. Primates, 35, 137–154.

27. Tomasello M and Camaioni L. 1997. A comparison of the gestural communication of apes and human infants. Human Development 40: 7–24.

